# The glycoprotease CpaA secreted by medically relevant *Acinetobacter* species targets multiple *O*-linked host glycoproteins

**DOI:** 10.1101/2020.07.22.216978

**Authors:** M. Florencia Haurat, Nichollas E. Scott, Gisela Di Venanzio, Juvenal Lopez, Benajmin Pluvinage, Alisdair B. Boraston, Michael J. Ferracane, Mario F. Feldman

## Abstract

Glycans decorate proteins and affect their biological function, including protection against proteolytic degradation. However, pathogenic, and commensal bacteria have evolved specific glycoproteases that overcome the steric impediment posed by carbohydrates, cleaving glycoproteins precisely at their glycosylation site(s). Medically relevant *Acinetobacter* strains employ their type II secretion system (T2SS) to secrete the glycoprotease CpaA, which contributes to virulence. Previously, CpaA was shown to cleave two *O*-linked glycoproteins, factors V and XII, leading to reduced blood coagulation. In this work, we show that CpaA cleaves a broader range of *O*-linked human glycoproteins, including several glycoproteins involved in complement activation, such as CD55 and CD46. However, only CD55 was removed from the cell surface, while CD46 remained unaltered during the *A. nosocomialis* infection assay. We show that CpaA has a unique consensus target sequence that consists of a glycosylated serine or threonine residue after a proline residue (P-S/T), and its activity is not affected by sialic acids. Molecular modeling and mutagenesis analysis of CpaA suggest that the indole ring of Trp493 and the ring of the Pro residue in the substrate form a key interaction that contributes to CpaA sequence selectivity. Similar bacterial glycoproteases have recently gained attention as tools for proteomic analysis of human glycoproteins, and CpaA appears to be a robust and attractive new component of the glycoproteomics toolbox. Combined, our work provides insight into the function and possible application of CpaA, a member of a widespread class of broad-spectrum bacterial glycoproteases involved in host-pathogen interactions.

**IMPORTANCE:** CpaA is a glycoprotease expressed by members of the *Acinetobacter baumannii-calcoaceticus* complex and it is the first *bona fide* secreted virulence factor identified in these species. Here, we show that CpaA cleaves multiple targets precisely at *O*-glycosylation sites preceded by a Pro residue. This feature, together with the observation that sialic acid does not impact CpaA activity, makes of this enzyme an attractive tool for the analysis of *O*-linked human protein for biotechnical and diagnostic purposes. Previous work identified proteins involved in blood coagulation as targets of CpaA. Our work broadens the set of targets of CpaA, pointing towards additional roles in bacteria-host interactions. We propose that CpaA belongs to an expanding class of functionally-defined glycoproteases that targets multiple *O*-linked host glycoproteins.

## INTRODUCTION

Members of the *Acinetobacter baumannii-calcoaceticus* complex are a frequent cause of serious multidrug resistant infections that are associated with high mortality, and they are a top priority for the research and development of new antimicrobial therapies (1, 2). However, the development of innovative therapies against these Gram-negative pathogens demands a better understanding of the virulence and resistance mechanisms critical to *Acinetobacter* infection. Previous work has implicated the type II secretion system (T2SS) in the pathogenesis of several *Acinetobacter* spp. in different infection models (3–8). The T2SS is a trans-envelope machine that mediates the transport of effector proteins from the periplasm to the extracellular milieu. First, T2SS effectors are translocated to the periplasmic space by either the general secretory pathway or the twin-arginine (TAT) system (5). Once in the periplasm, the effectors fold into their tertiary/quaternary structure, a process occasionally facilitated by dedicated chaperones, and are subsequently targeted to the T2SS machinery for secretion (7). Although the T2SS machinery is conserved across *Acinetobacter* spp, the effector repertoire is diverse and varies from strain to strain (7, 9, 10)

Approximately 20 to 60 T2SS effectors are secreted by any given *Acinetobacter* strain, and these proteins are involved in lipid assimilation, serum resistance, colonization of various host tissues, antibiotic resistance, and biofilm formation (3, 4, 7, 11). Among these T2SS effectors, the zinc-metallo-endopeptidase CpaA is the best-characterized and most abundant effector secreted by several medically relevant *Acinetobacter strains* (7, 8, 12). We demonstrated that CpaA stability and secretion depends on the membrane-bound chaperone CpaB, which is encoded adjacent to CpaA (7, 8). The N-terminal transmembrane domain of CpaB anchors the protein to the membrane, whereas its C-terminal domain interacts directly with CpaA in the periplasm (8). Removal of the CpaB transmembrane domain results in secretion of the CpaA-CpaB complex (8). Structural studies confirmed that CpaA and CpaB strongly interact in a 1:1 ratio via a novel protease-chaperone arrangement in which CpaA surrounds CpaB (8, 13). This unusual configuration was not observed in other previously characterized T2SS chaperone/effector pairs (14). An additional interesting feature observed in the co-crystal structure is that the CpaB C-terminal tail blocks access to the CpaA catalytic site (13).

In *A. nosocomialis* M2, deletion of *cpaA* has comparable virulence defects to those observed in a T2SS mutant strain (8). Importantly, using a murine pneumonia model, we demonstrated that CpaA plays a crucial role in dissemination to the spleen (8). The link between CpaA and dissemination within the host is further supported by *in vitro* data showing that CpaA is able to cleave human factor V (fV) and factor XII (fXII), thus interfering with blood coagulation (12, 15). Interestingly, CpaA cleaves fXII at two *O*-glycosylation sites in its proline-rich region, between Pro_279_Thr_280_ and Pro_308_Thr_309_, with both Thr residues being *O*-glycosylated, though glycans are known to protect proteins from degradation (15). Moreover, glycosylation is required for CpaA activity, as deglycosylation of fXII abrogates CpaA activity (15). However, the full scope of CpaA substrates and the basis for their recognition by CpaA remain poorly understood.

Few glycoprotein-targeting zinc-metallo-endopeptidases have been structurally characterized. They belong to either the metzincin or gluzincin protease superfamilies (16, 17). Metzincin proteases have a characteristic extended zinc-binding motif (HExxHxxGxxH) and a conserved Met-turn, whereas gluzincin proteases have a glutamate residue in the zinc-binding motif (HExxHE) (18). We previously determined the X-ray crystal structure of CpaA (13). Key features of its catalytic domain (α+β fold, extended zinc-binding motif and a conserved Met-turn) revealed that CpaA belongs to the metzincin protease superfamily (13, 19). In addition to its catalytic domain, CpaA possesses four very similar β-sheet tandem repeats, all of which share a similar Ig-like folds and resemble glycan-binding domains (13, 15). Interestingly, these repeats share fold similarities to a domain present in the metzincin protease StcE, a secreted glycoprotease from Enterohemorrhagic *Escherichia coli* (EHEC), though their detailed sequence and structural similarities are very limited (13, 20). StcE is also a T2SS secreted protease known to target highly *O*-glycosylated epithelial substrates, such as CD55 (16, 17, 21–23).

Given these reported structural similarities, we hypothesized that CpaA could target a broader number of glycosylated host proteins. In this work, we combine biochemistry, mass spectrometry (MS), molecular modeling and *in vivo* assays to demonstrate that CpaA targets multiple *O*-linked human glycoproteins. This work furthers our understanding of the interaction of CpaA with its targets and provides valuable insight into *Acinetobacter* pathobiology. Furthermore, we show that CpaA has a possible biotechnological application as a tool for glycoproteomics.

## RESULTS

### CpaA is a broad-spectrum glycoprotease

Previous reports indicated that CpaA targets coagulation-related glycoproteins, such as fV and fXII (12, 15). CpaA cleaves the fXII mucin-like region at two sites (15). Mucins are a family of high molecular weight glycoproteins composed of Pro/Thr/Ser-rich domains, that are heavily decorated with long oligosaccharides (24, 25). While glycosylation is protein-dependent, the average mucin is ~50% *O*-linked glycan by mass, and these glycans tend to be very heterogeneous as results of unique extensions and branching of the core structure (24, 25). Glycoproteases targeting mucins showcase a wide variability in terms of substrate preferences, even among members of the same family, and therefore prediction of their targets is not possible (16, 21–23, 26–29). To assess whether CpaA can cleave other glycoproteins, we expressed and purified CpaA and its catalytically inactive point mutant (E520A) as C-terminal His-tagged proteins in *A. nosocomialis* M2, as previously done (8, 13). CpaA and CpaA_E520A_ were incubated overnight with several medically relevant and commercially available *O*-linked glycoproteins (CD55, CD46, TIM1, TIM4 and C1-INH). Mucins run as large and diffuse bands as a result of their heterogeneity in size and glycosylation. Remarkably, CpaA, but not catalytically inactive CpaA_E520A_, cleaved all the tested mucin glycoproteins, as observed by gel shifts to lower molecular weight (**Figure 1A**, degradation products are indicated with asterisks). We also tested the ability of CpaA to cleave two well-characterized recombinant therapeutic glycoproteins, etanercept (Enbrel) and abatacept (Orencia). These fusion proteins are highly glycosylated proteins purified from mammalian cells and contain mucin-like domains in their structures (30–32). Once again, CpaA, but not CpaA_E520A_, cleaved both recombinant glycoproteins, as observed in **Figure 1B** (degradation products are indicated with asterisks).

**Figure 1.**
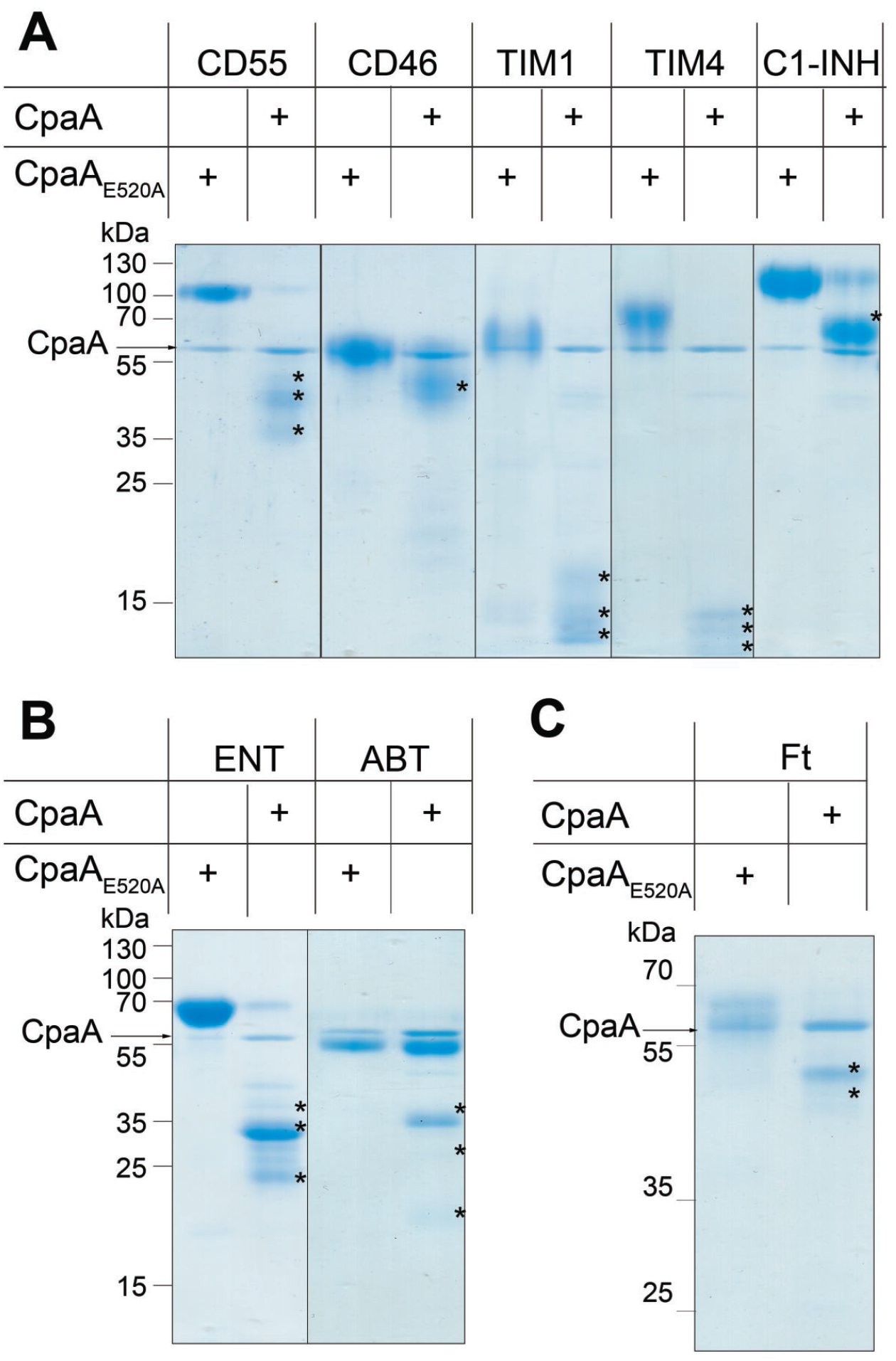
CpaA targets multiple proteins in vitro. Purified **(A)** mucins, **(B)** mucin-like proteins and **(C)** O-glycoproteins were treated with purified CpaA and catalytically inactive CpaA (CpaA_E520A_) (~55kDa). Samples were separated by SDS-PAGE. Digestion products are indicated by an asterisk. These data are representative of at least 3 independent experiments C1-INH: C1 esterase inhibitor. TIM: T-cell immunoglobulin and mucin domain. ENT: etanercept. ABT: abatacept. Ft: Bovine fetuin

### CpaA specifically targets O-linked glycoproteins and its activity is unaffected by sialic acid

The activity of glycoproteases is differentially impacted by many factors, such as *O*-glycosylation density and glycan composition. For instance, StcE and TagA, a secreted glycoprotease from *Vibrio cholerae*, uniquely cleave heavily glycosylated proteins, such as mucins, but they cannot digest less *O*-glycosylated proteins such as fetuin (Ft). Fetuin, also known as alpha-2-HS-glycoprotein, is decorated with up to five *O*-linked sialylated glycans, and is often used as a model protein for glycosylation-related studies (22, 26, 33). In contrast, the activity of other glycoproteases, such as the gluzincin IMPa from *Pseudomonas aeruginosa,* is unaffected by the degree *O*-glycosylation (16). Like IMPa, CpaA cleaved fetuin (**Figure 1C**), but not enzymatically deglycosylated fetuin (**Figure 2A**). The activity of various glycoproteases can be dependent on, inhibited by, or unaffected by the presence of sialic acid, a negatively charged 9-C sugar with key biological functions (34). For example, glycoproteases from *Mannheimia haemolytica* and *Clostridium perfringens* require sialic acid for activity, whereas other glycoproteases are inhibited by this sugar (16, 35, 36). StcE is an example of a glycoprotease that is indifferent to sialic acid (22). Similarly to StcE, CpaA efficiently digested asialo-fetuin (AFt), a variant of fetuin lacking only the sialic acid moieties (**Figure 2B**), indicating that CpaA glycoprotease activity is sialic acid-independent.

**Figure 2.**
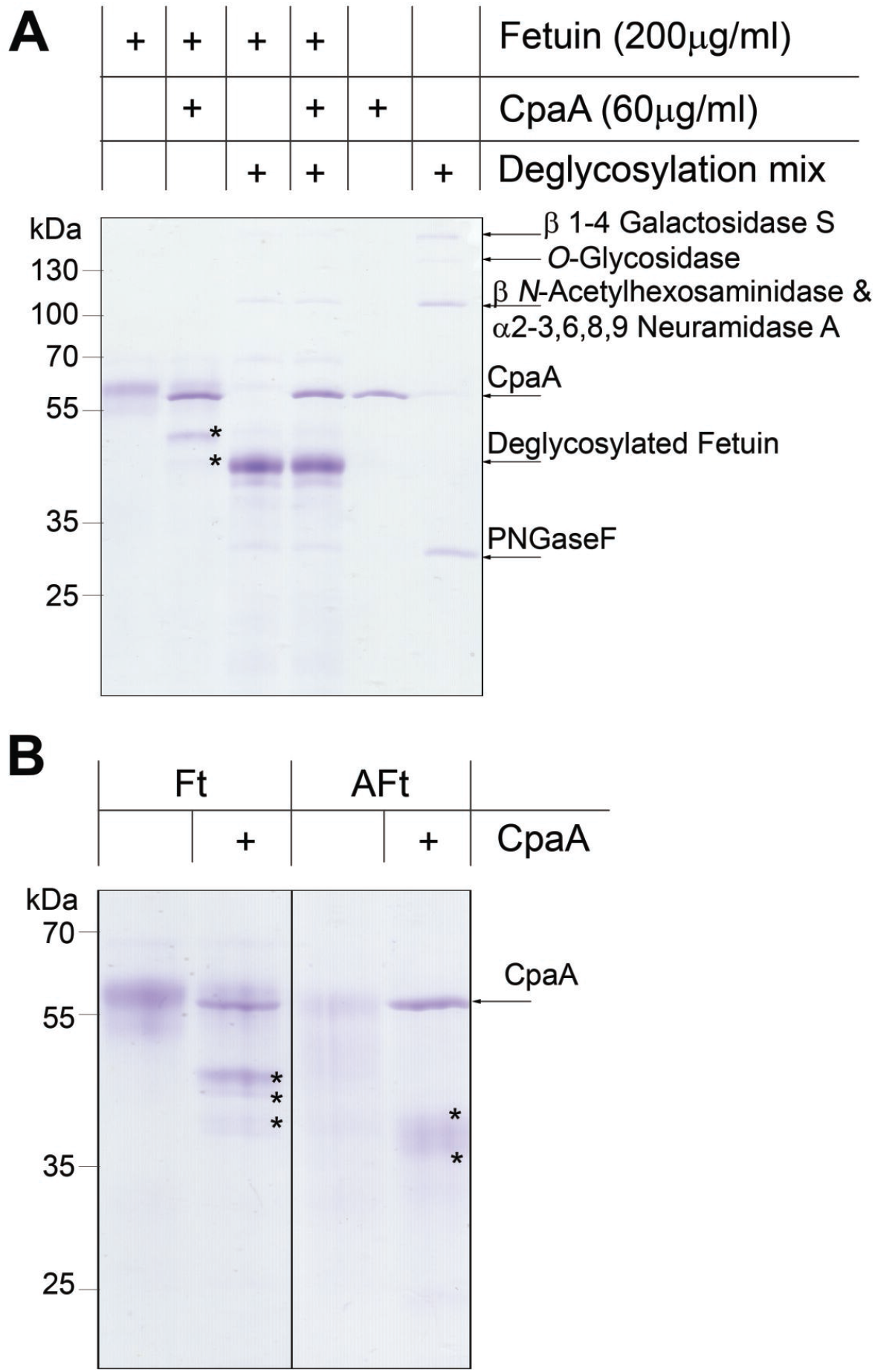
Fetuin glycosylation is required for CpaA activity. **(A)** Fetuin and fetuin treated with a deglycosylation mix were incubated with CpaA. Digestion products (indicated with an asterisk) are only observed for fetuin. **(B)** The presence of sialic acid does not affect CpaA activity. Ft: Bovine fetuin. Aft: asialofetuin.

All of the CpaA targets tested so far were simultaneously *N*- and *O*-glycosylated proteins. To test whether CpaA can cleave *N*-linked glycoproteins, we incubated CpaA with RNAseB, a glycoprotein modified only with *N*-linked glycans. No proteolytic activity was observed with RNAseB as a substrate (**Figure S1**). Together, these results indicate that CpaA possesses broad substrate specificity, being able to cleave multiple *O*-glycosylated proteins in addition to fV and fXII, and that CpaA activity is unaffected by sialylation.

### CpaA cleaves between Pro and a glycosylated Ser/Thr

We employed a MS approach to gain insight into the common molecular features that dictate CpaA substrate recognition. We excised the gel pieces containing the proteolysis products indicated with asterisks in **Figure 1**, treated them with trypsin and subsequently subjected them to MS analysis, as done previously (37). For most of the proteins, with the exceptions of TIM4 and C1-INH, we identified non-tryptic peptides, resulting from CpaA activity (**Figure 3** and **Figure S2**). These peptides contained an invariant C-terminal Pro residue (P1 position) (**Figure 3A** and **Figure S2A, C and D**), which in the full-length protein sequences, is always adjacent to a glycosylated Ser or Thr (S/T*) (**Figure 3C**) We also identified non-tryptic glycopeptides derived from etanercept that contained a glycosylated N-terminal Ser (**Figure 3B** and **Figure S2B**). The peptides were glycosylated with either *N*-acetyl hexose-hexose-*N*-acetyl neuraminic acid (HexNAc-Hex-Neu) or HexNAc-Hex moieties (**Figure 3B** and **Figure S2B**), reinforcing the concept that CpaA activity is indifferent to the presence of sialic acid. The CpaA-dependent cleavage products, including those of fXII (previously reported), were used as WebLogo inputs (**Figure 3C**) (15, 38). This analysis revealed that CpaA has a distinct peptide consensus sequence, P-S/T*, where cleavage occurred before the glycosylated Ser/Thr (S/T*) residue. As seen in **Figure 3C**, the C-terminal Pro residue preceding the *O*-glycosylation site is likely a strict requirement for CpaA targeting. Human erythropoietin (EPO) contains the same Ser-bound oligosaccharide as fetuin but has an Ala residue preceding the glycosylation site. This protein was not cleaved by CpaA (**Figure S3**), further supporting the essentiality of the Pro residue preceding the *O*-glycosylation site.

**Figure 3.**
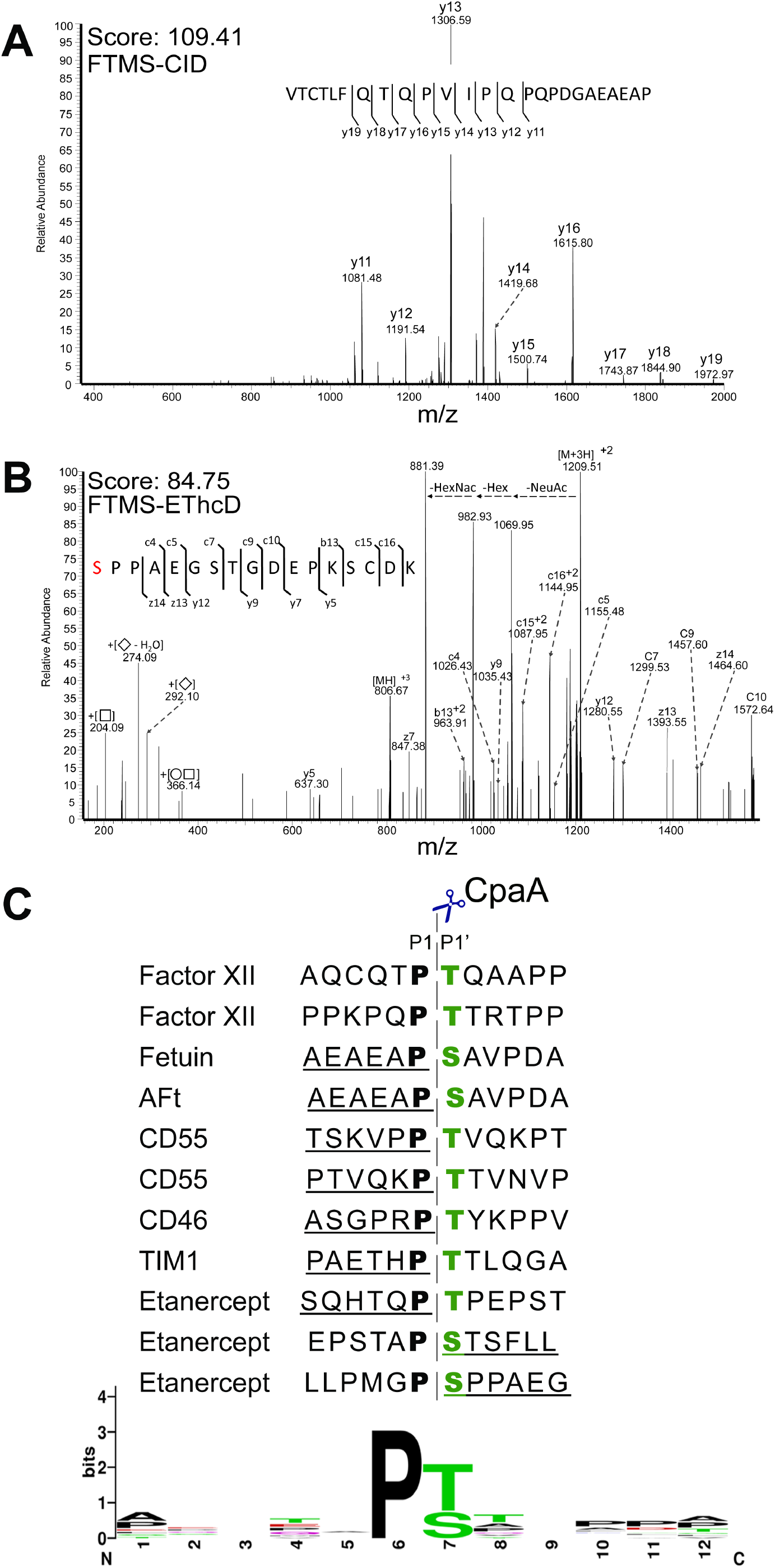
CpaA cleaves between Pro and a glycosylated Ser/Thr. Mass spectra of the non-tryptic fragment of **(A)** Fetuin and **(B)** Entanercept. The glycosylated Ser, residue, indicated in, red, is modified with HexNAcHexNeuAc. **(C)** Sequence of the CpaA-dependent cleavage products were used as WebLogo inputs (weblogo.berkeley.edu). Factor XII cleavage sites were previously reported (15). The underlined peptide sequences were detected by mass spectrometry. The dashed line indicates CpaA cleavage site. HexNAc: *N*-acetyl hexosamine. Hex: hexose. NeuAc: *N*-acetyl neuraminic acid.

### Mammalian glycan array screening

The recombinant proteins tested in this study were all expressed and purified from HEK293 cells. This cell line generates proteins *O*-glycosylated with disaccharide structures galactose β1-3 *N*-acetylgalactosamine-(Galβ1-3GalNAc-), also known as Core I, as well as mono- and di-sialylated core I and core II (Galβ1-3[GlcNAcβ1-6]GalNAc-) structures (39). Other glycan cores have more complex (branched) structures that may or may not be accommodated by CpaA (40). To this end, we employed a mammalian glycan array screening assay (Consortium for Functional Glycomics) to test if CpaA can directly bind various glycans. However, we did not detect any significant peak indicative of binding for any glycan in the array when employing two protein concentrations (5 μg/ml and 50 μg/ml) of either CpaA or its catalytically inactive mutant CpaA_E520A_, (**Figure S4A and S4B**). A similar lack of binding has been previously reported for other glycoproteases, which may be due to low-affinity or transient interactions with the glycans (41). It is noteworthy that none of the glycans in the glycan array are conjugated to P-S/T sequence. Considering our previous results, it is likely that recognition by CpaA is dependent on a combination of both protein sequence and glycan structure adopted in the array.

### CD55 is removed from epithelial surfaces by secreted CpaA

Our previous *in vitro* results expanded the known CpaA substrates to include human glycoproteins beyond those involved in blood coagulation (**Figure 1A**). To gain insight about CpaA activity in the context of an infection, we tested whether CpaA directly cleaves surface exposed *O*-glycoproteins. Of all the glycoproteins tested *in vitro,* CD55 and CD46 are highly expressed cell surface *O*-glycoproteins in HeLa cells(42). Thus, HeLa cells were treated with purified CpaA and CpaA_E520A_, and the levels of CD55 and CD46 bound to the cell surface were quantified by flow cytometry (**Figure 4**). At the two protein concentrations tested, cells treated with CpaA displayed a reduced amount of CD55 on their surface compared to those treated with CpaA_E520A_ (**Figure 4A**). In contrast, although CpaA was able to cleave CD46 in our *in vitro* assay (**Figure 1A**), cell surface exposed CD46 levels remained unaltered after CpaA treatment, independent of the protein concentration used (**Figure 4B**).

**Figure 4.**
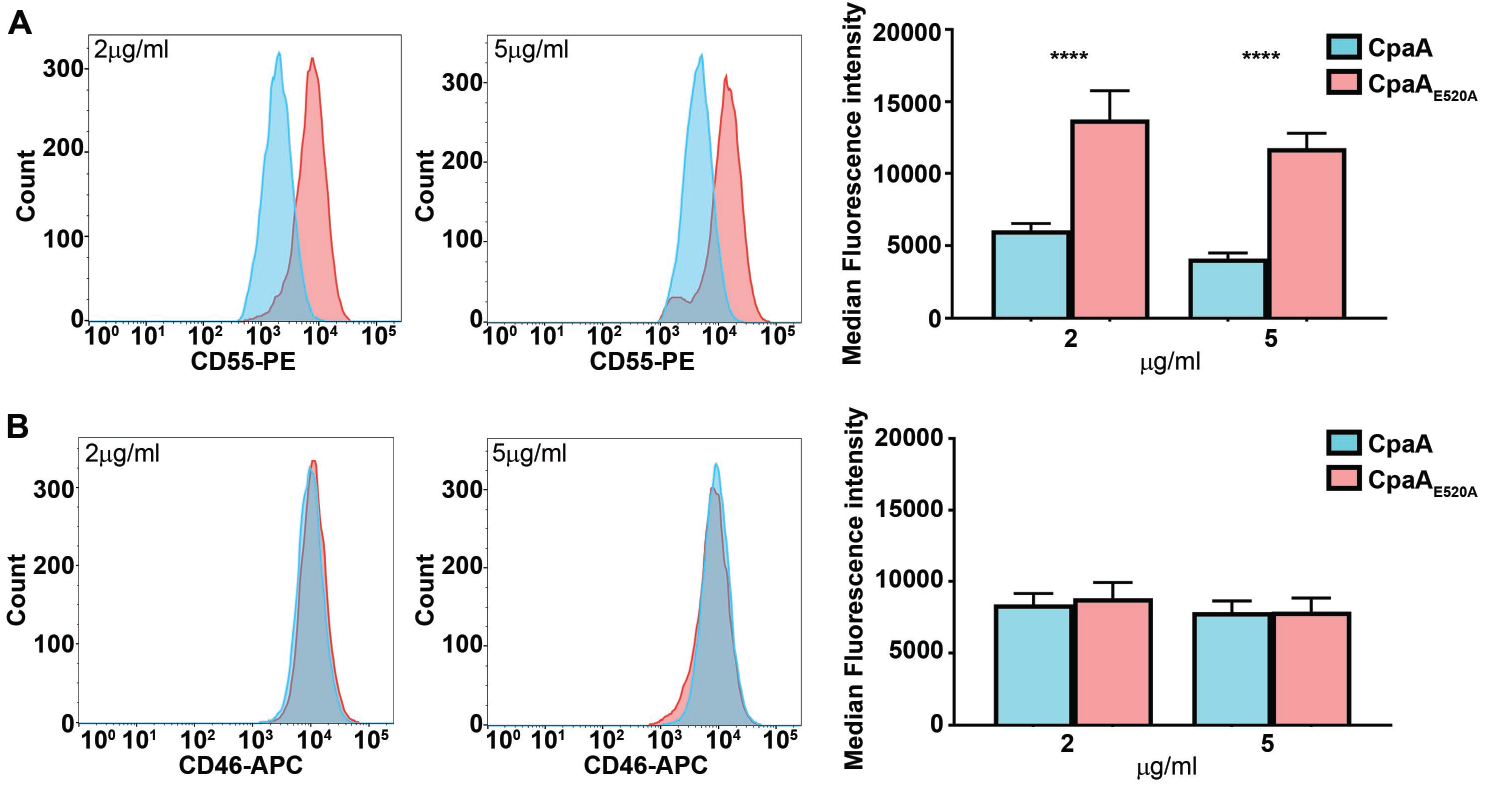
Purified CpaA cleaves CD55 but not CD46 from HeLa cell surface. Surface level of **(A)** CD55 and **(B)** CD46 were measured by flow cytometry on HeLa cells. Cells were incubated with two different concentrations (2 μg/ml and 5 μg/ml) of purified CpaA and CpaA_E520A_ for 1h. Left and middle panels are representative flow cytometry histograms. In the right, panel, median ± S.D of three biological repeats are shown. ***, *p* < 0.0001 Two-Way ANOVA with Sidak’s multiple comparisons test.

Next, we infected HeLa cells with *A. nosocomialis* M2 expressing CpaA and CpaA_E520A_ at three different MOI (10, 100 and 1000), and used flow cytometry analysis to quantify the levels of cell surface exposed CD55 and CD46 post-infection (**Figure 5**). CD55 and CD46 levels remained unchanged when cells were infected with *A. nosocomialis* M2 secreting CpaA_E520A_, indicating that *A. nosocomialis* M2 does not secrete any other proteases targeting either glycoprotein (**Figure 5**). In agreement with our previous results, secreted CpaA cleaved CD55 but not CD46 from the cell surface (**Figure 5**). The CD46 protein used in the *in vitro* assay (**Figure 1A**) was expressed and purified from HEK293 cells. It is well known that glycosylation patterns differ between cell lines (43), thus, it is possible that different protein glycosylation patterns impact CpaA activity. To address this, we repeated the experiment infecting HEK293 cells. As observed with HeLa cells, secreted CpaA digested CD55 but not CD46 (**Figure S5**), indicating that potential differences in glycosylation between these two cells lines does not account for these discrepancies. Importantly, a MOI of 100 was sufficient to detect cleavage of CD55 from HeLa cells and increasing the MOI to 1000 did not boost CD55 cleavage by CpaA (**Figure 5A**). The remaining CD55 (and perhaps CD46) may be associated with proteins/ligands that prevent CpaA activity. We conclude that *A. nosocomialis* secretes physiological levels of CpaA that can digest host surface exposed proteins during infection.

**Figure 5.**
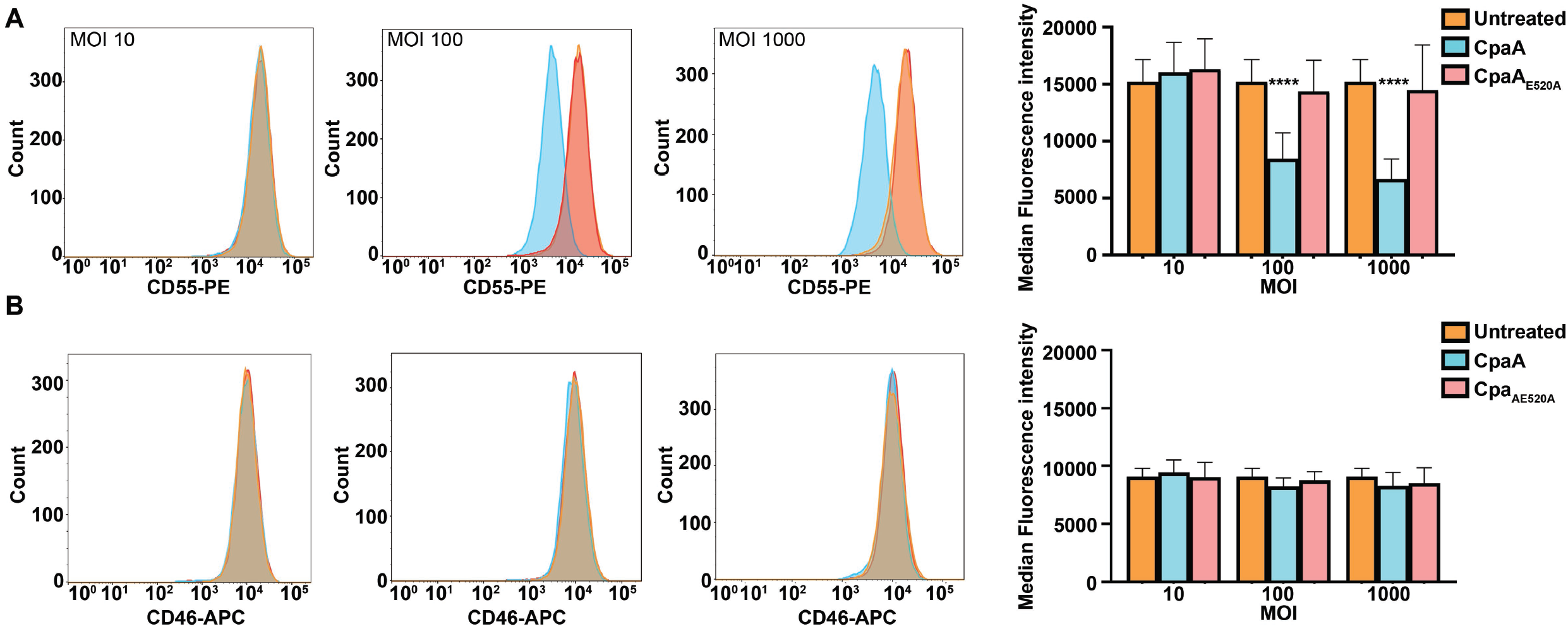
Secreted CpaA cleaves CD55 from HeLa cell surfaces. Surface level of **(A)** CD55 and **(B)** CD46 were measured by flow cytometry on HeLa cells. Cells were incubated with three different MOI (10, 100, 1000) of *A. nosocomialis* M2 secreting either CpaA or CpaA_E520A_ for 2.5h. Panels are representative flow cytometry histograms. In the right panel, median ± S.D of three biological repeats are shown. ***, *p* < 0.0001 Two-way ANOVA with Dunnett’s multiple comparisons test.

### Molecular modeling identifies putative binding mode of glycopeptides to CpaA

We previously determined the X-ray co-crystal structure of the CpaA-CpaB complex (13), and others have obtained crystal structures of related metzincin enzymes with peptide- and peptidomimetic-based ligands (**Table 1**) (44, 45). X-ray structures of the related gluzincin glycopeptidases BT4244 (*Bacteroides thetaiotaomicron*), IMPa, and ZmpB (*Clostridium perfringens* ATCC 13124) have also been determined with bound glycoamino-acid and glycopeptide ligands (16). Notably, these gluzincin enzymes cleave fetuin, asialofetuin, and related synthetic glycopeptides to varying degrees (**Table 1**). Thus, to better understand binding between CpaA and glycopeptide substrates, we performed docking experiments between a CpaA model and three different glycoforms of a fetuin-based fragment peptide (Ac-EAPSA-*N*Me, where S is glycosylated). One of the glycoforms lacks the sialic acid moiety (**Figure 6** and **Figure S6A**), while the other two are sialylated at two different position of the Galβ1-3GalNAc-core (**Figure S6B-C**). The peptide portions of the docked species were able to contact the catalytic zinc ion while forming an antiparallel β-sheet with a β-strand of the active site (**Figure 6A**), findings that are consistent with aforementioned metzincin crystal structures as well as docking experiments with StcE (16, 22). In the docked structures, the consensus P1 proline residue lies adjacent to W493 and is somewhat solvent-exposed (**Figure 6B**). An H-pi interaction was observed between a proline beta hydrogen and the tryptophan indole ring (**Figure 6B**). These studies suggest that CpaA selectivity may be the result of W493 (a) forming a potential H-pi interaction with the prolyl ring, (b) minimizing the prolyl residue’s exposure to solvent and/or (c) sterically holding the substrate in the active site.

**Figure 6.**
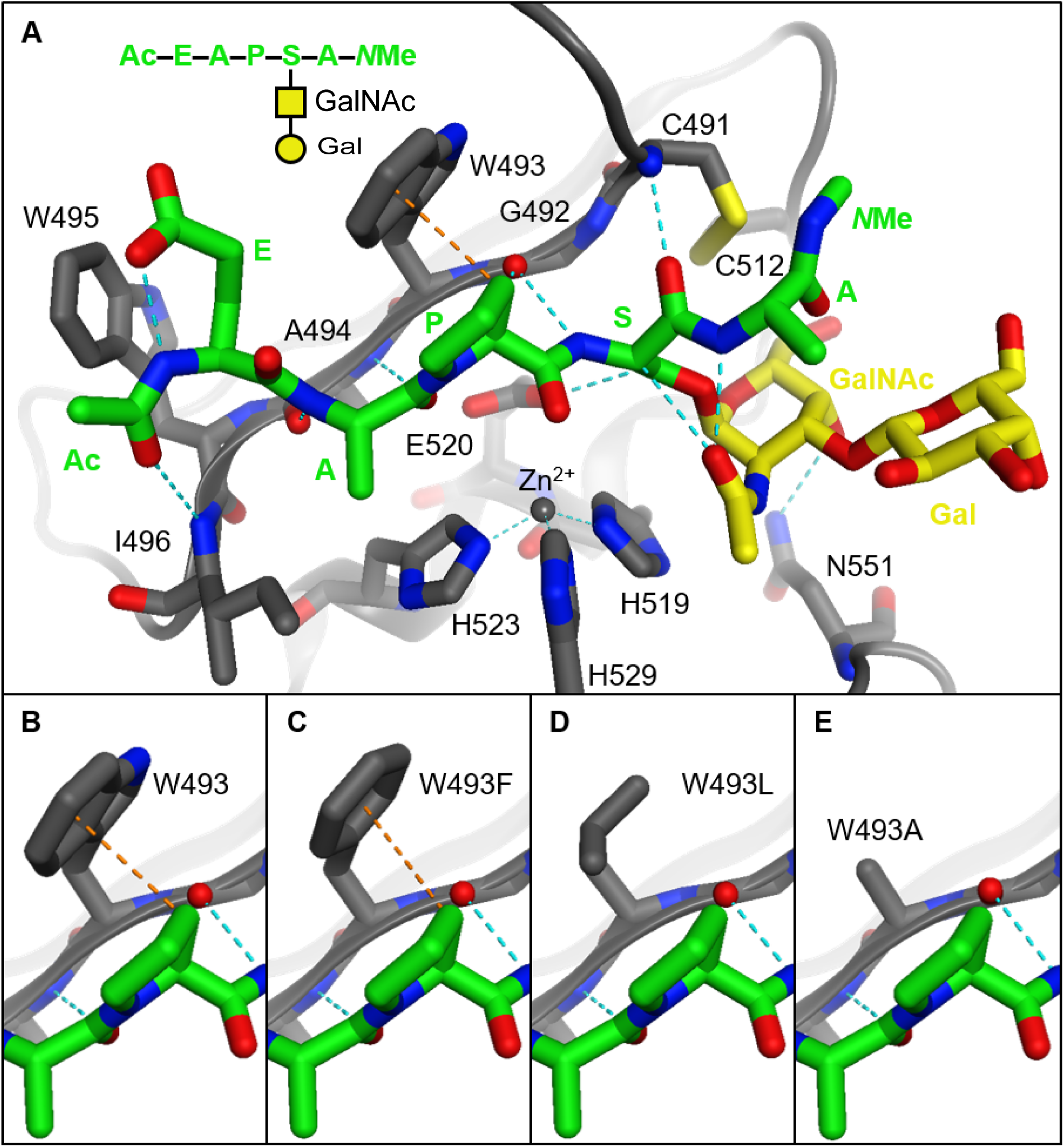
Structure of a fetuin-based model glycopeptide substrate docked in models of CpaA and related mutants. **(A)** The enzyme (gray) and the substrate’s peptide portion (green) form hydrogen bonds (dashed blue lines) between their backbones similar to an antiparallel beta sheet. The catalytic histidine residues (gray sticks) and zinc ion (gray sphere) hydrolyze the amide bond between the substrate’s proline (P1) and glycosylated serine (P1’) residues. The GalNAc moiety (yellow sticks) forms hydrogen bonds with the substrate and the enzyme, whereas the Gal moiety (yellow sticks) is exposed to solvent. **(B-E)** The tryptophan residue of the native enzyme can form an H-pi interaction (dashed orange lines) with the substrate. This is interaction is weakened when this residue is mutated to phenylalanine and nonexistent with aliphatic residues.

**Table 1.** Summary of the biochemical and structural features of known *O*-glycoproteases.

| Glycoprotease | Source | Secretion | MEROPS | Family | Structural domain organization | Target Mucins | Target Fetuin | Sialic acid requirement | Recognition site | Ref |
| --- | --- | --- | --- | --- | --- | --- | --- | --- | --- | --- |
| CpaA | *A* spp. | T2SS | M72 | Metzincin | IG 1 IG 2 IG 3 IG 4 CD | Yes | Yes | Indifferent | P-S/T* | This study and (13) |
| StcE | EHEC | T2SS | M66 | Metzincin | IG INS IG M-SD1 M-SD2 D1 D2 CD | Yes | No | Indifferent | S/T*X-S/T | (17, 22, 33) |
| BT_4244 | *Bt* | Extracellular Lipoprotein | M60 | Gluzincin | IG 1 CD IG 2 | Yes | No | No sialic acid | T/S-T/S* | (16) |
| IMPa | *Pa* | T2SS | M88 | Gluzincin | $\alpha/\beta$ H IG CD | Yes | Yes | Its removal enhances activity | N.D. | (12, 67, 68) |
| ZmpB | *Cf* | SP | M60 | Gluzincin | IG 1 CD IG 2 | Yes | Yes | Required | TE-T/ST/S | (16) |
| Pic | *Ec* | T5SS | S6 | SPATE | Structure undetermined | Yes | No | Required | T/S-S/T* | (51, 69, 70) |
| ZmpC | *Sp* | Sortase | M26 |  | Structure undetermined | Yes | Yes | Indifferent | T*XXX-X | (51) |
| AMUC_0627 | *Am* | SP, Extracellular unknown | M60 | Gluzincin | Structure undetermined | Yes | No | Its removal enhances activity | S/T*-S/T* | (51) |
| AMUC_0908 | *Am* | SP, Extracellular unknown | M60 | Gluzincin | Structure undetermined | Yes | No | Indifferent | T/S-T/S* | (51) |
| AMUC_1514 | *Am* | Extracellular unknown | M60 | Gluzincin | Structure undetermined | Yes | No | No sialic acid | T/S-T/S* | (51) |
| TagA | *Vc* | T2SS | M66 | Metzincin | Structure undetermined | Yes | No | N.D. | TE-T/ST/S | (51) |
| SslE/YghJ | *Ec/Sf* | T2SS | M98 | Gluzincin | Structure undetermined | Yes | N.D. | N.D. | N.D. | (28, 71) |
| O-sialoglycoprotease | *Mh* | Extracellular, unknown | M22 | N.A. | Structure undetermined | Yes | No | Required | N.D. | (70, 72) |

N.D. Not determined. N.A.: Not applicable. CD: Catalytic domain. IG: Ig-like domain. INS: Insertion domain. M-SD: metalloprotease module. D: disordered region. $\alpha/\beta$ : $\alpha/\beta$ domain. H: helix bundle linker *Bt*: *Bacteroides thetaiotaomicron*. *Pa*: *Pseudomonas aeruginosa*. *Cf*: *Clostridium perfringens*. *Ec*: *Escherichia coli*. *Am*: *Akkermansia muciniphila*. *Vc*: *Vibrio Cholerae*. *Sf*: *Shigella flexneri*. *Mh*: *Mannheimia haemolytica*

In the docked structures, the glycan moieties were found to form few interactions with CpaA and the substrate peptide. The acetyl group of the GalNAc moieties all formed hydrogen bonds with the peptide backbone (**Figure 6A**). Similar contacts were observed in docking studies using *O*-GalNAcylated peptides and StcE (22). Such contacts have been shown to affect the conformation of mucin-like glycopeptides (46), and they may bias substrates into an extended conformation that would be more easily recognized by the enzyme. We also found that the 4-OH of the GalNAc moieties formed hydrogen bonds with the side chain amide of N551 (**Figure 6A** and **Figure S6A-C**). This interaction would likely impart the enzyme with selectivity for peptides modified with GalNAc at P1’. Interestingly, the side chain of this residue occupies similar space as the side chains of tryptophan residues that are conserved in the related gluzincin enzymes BT4244, IMPa, and ZmpB (**Figure S6D-E**). In the gluzincin enzymes, the indole nitrogens instead hydrogen bond to the acetyl groups of GalNAc ligands. Such a contact may be formed between CpaA and its substrates if minor conformational changes occur. In either case, N551 appears to be the main residue recognizing the unique features of the GalNAc moiety, as the other residues flanking the glycans are mostly small and nonpolar. The related gluzincin enzymes, on the other hand, form several contacts between polar side chains and their GalNAc groups.

Larger, linear glycans project subsequent sugar residues into solvent; these groups are not predicted to interact with CpaA. Branched glycans are not well accommodated by the enzyme, as the disulfide bond adjacent to the active site limits the flexibility of the enzyme and the branched sugar’s ability to bind. Conversely, both IMPa and ZmpB form interactions with additional sugars of their corresponding glycans. Still, only IMPa demonstrated binding activity in the mammalian glycan array screen (16), which may explain why no hits were found in the same screen with CpaA. Collectively, our docking studies indicate that CpaA interacts with both peptide and glycan components of the substrate and identify residue W493 as a potential mediator of the interaction between CpaA and the Pro residue of its substrate.

### Effect of W493 on CpaA specific activity

Docking studies suggest an H-pi interaction between the indole ring (of W493) of CpaA and a beta hydrogen of Pro residue (at the P1 position) of the substrate (**Figure 6B**). While other substrate residues can form this contact with W493, the unique rigidity and geometry of Pro allows it to optimally present its beta hydrogen to the indole ring of W493. Thus, we hypothesized that W493 of CpaA plays a critical role in CpaA selectivity. Indeed, our molecular docking studies indicate a weaker H-pi interaction when peptides were docked into a mutant W493F model (**Figure 6C**). The modelled W493L and W493A mutants have aliphatic residues that are unable to form this interaction with the substrate (**Figure 6D** and **Figure 6E**). Moreover, all the mutant models formed fewer van der Waals contacts with the Pro residue and further exposed it to solvent.

To complement our molecular modeling experiments, we tested the effect of these mutations on CpaA activity. All CpaA point mutation variants were expressed and secreted at similar levels to CpaA and CpaA_E520A_, and no degradation products of CpaA were observed in the whole cell fractions of the CpaA variants (**Figure S7A and B**). CpaA_E520A_ was included as negative control for CpaA activity. We purified all His-tagged CpaA variants and determined their *in vitro* activity against various substrates (**Figure S7C**, **Figure 7A** and **Figure S8**). The different mutations affected CpaA efficiency and site recognition in a substrate-specific manner. All mutants, except for CpaA_E520A_, were able to cleave fetuin, yielding a similar cleavage pattern (**Figure 7A**). However, CpaAW493A cleaved fetuin with less efficiency. None of the CpaA variants were able to cleave EPO, further highlighting the essentiality of a Pro residue at P1 for targeting by CpaA (**Figure 7B**). Treatment of CD55 and C1-INH with the CpaA mutants revealed that all variants are less active, shown by an increase in the amounts of undigested substrate. Additional faint bands were observed, but we were unable to determine the cleavage site using MS analysis (**Figure S8**).

**Figure 7.**
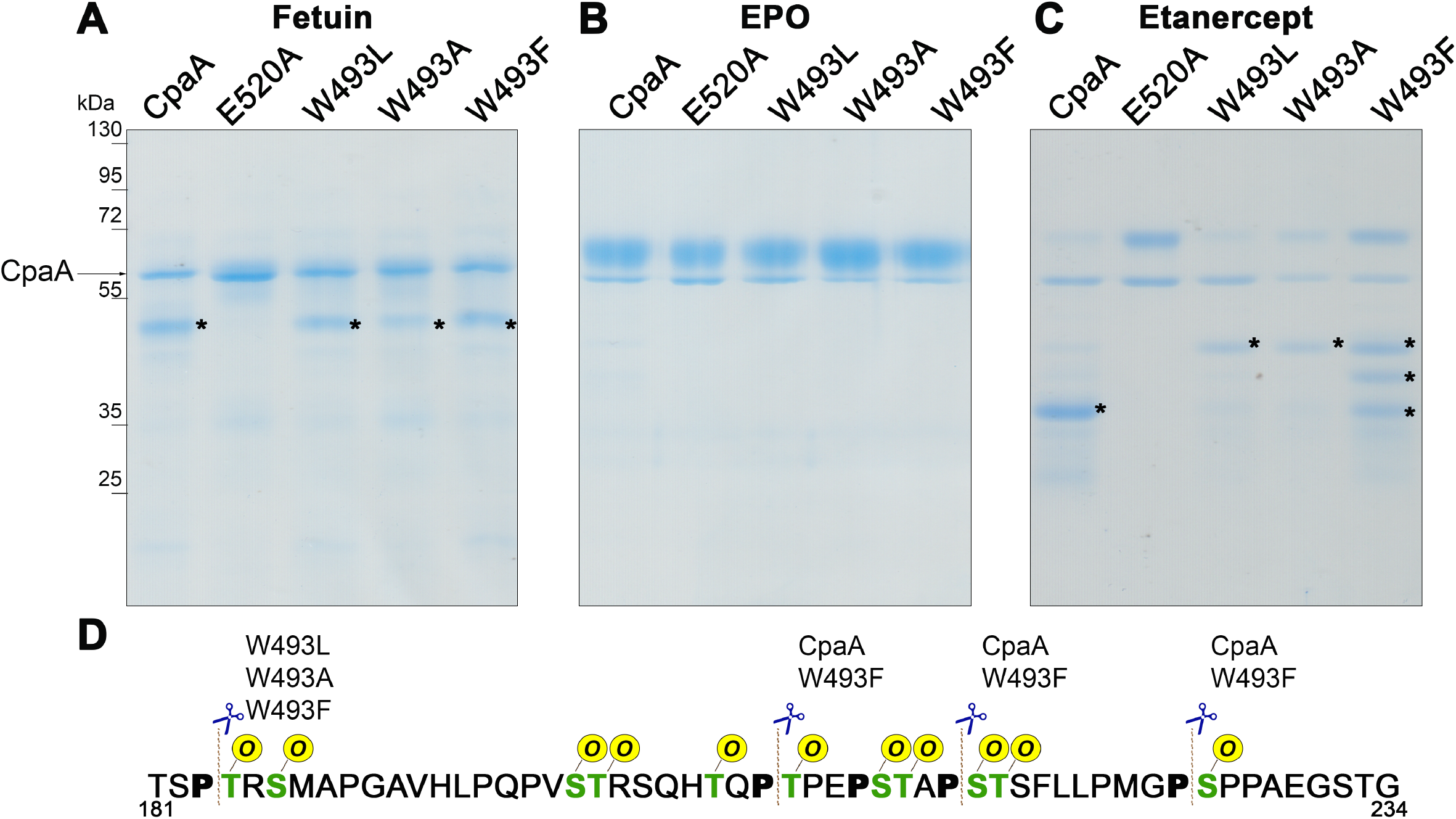
W493 affects CpaA activity. **(A)** Fetuin, **(B)** EPO and **(C)** Etanercept were incubated with purified CpaA and the different CpaA mutants (~55kDa). Samples were separated by SDS-PAGE stained with InstantBlue. Digestion products are indicated by an asterisk. These data are representative of at least 3 independent experiments. **(D)** Fragment of Etanercept protein sequence. MS analysis of proteins bands allowed the identification of the sites preferentially cleaved by the four CpaA variants. In green are all the known *O*-glycosylation sites and, in bold, the P next to the glycosites. The dashed line indicates the cleavage site of CpaA and the different point mutants.

We previously identified three cleavage sites for CpaA on the mid-region of etanercept, Pro_207_Thr_208_, Pro_215_Ser_216_ and Pro_225_Ser_226_ (**Figure 3B and S2**). Cleavage by CpaA generates two fragments of similar molecular weight that co-migrate as a single band of about ~36 kDa in SDS-PAGE (**Figure 1A** and **Figure 7C**). Notably, digestion of etanercept by CpaAW493A and W493L generated a major product of ~45 kDa instead (**Figure 7C**). CpaAW493F produced three bands that migrate as ~36, 40 and 45 kDa fragments respectively (**Figure 7C**). MS analysis of these bands allowed the identification of the sites preferentially cleaved by the three CpaA variants **(Figure 7D**). The unique ~45 kDa band results from cleavage at the Pro_183_Thr_184_ site, which is not a preferred site for wild-type CpaA. Unlike the other cleavage sites, the Pro_183_Thr_184_ site is in an area of low glycosylation of etanercept. Interestingly CpaAW493A and W493L variants were unable to cleave etanercept in the high glycosylation density region, which could explain the low activity against other mucin targets. Taken together, our data suggest that W493 plays a role in CpaA substrate selectivity by interacting with the Pro residue of its target protein.

### CpaA belongs to an expanding functionally-defined class of surface-exposed and secreted glycoproteases

Mounting evidence indicates that both commensal and pathogenic species produce and secrete glycoproteases to modulate adherence, penetrate the inner mucus layer, or evade the host immune response (18, 21, 47). For example, StcE contributes to immune evasion during EHEC infection by preventing immune cells from moving to the sites of infection (20, 23, 48, 49). Additionally, StcE activity against mucins promotes access of EHEC to epithelial cells which assists during host cell colonization (20, 50). *P. aeruginosa* produces IMPa, which cleaves the macrophage surface protein CD44 inhibiting phagocytosis (29). IMPa also cleaves P-Selectin Glycoprotein Ligand 1 (PSGL-1), helping the bacterium to escape neutrophil attack (29). In *Vibrio cholerae*, secreted TagA targets host cell surface glycoproteins, modulating bacterial attachment during infection (26). These examples are indicative of the pivotal role of glycoproteases in modulating host-pathogen interactions by targeting various host proteins. Despite their relevance, only a relatively small number of bacterial glycoproteases have been biochemically characterized to varying extents **(Table 1)**. These enzymes are commonly encoded by bacteria isolated from mucin-rich environments, such the human gut or lungs. These glycoproteases differ in their secretion mechanisms, protease class, domain organization, catalytic site, and recognized targets. Like IMPa, StcE, TagA, and SslE/ YghJ, CpaA is secreted by a T2SS. CpaA, IMPa, and ZmpC display broad *O*-glycoprotease activity targeting mucins, regardless of their glycosylation density, as well as *O*-glycoproteins with low *O*-glycan chain density (fetuin). On the other hand, StcE can cleave only mucins with long mucin-like regions (such as CD43, and CD55) or short mucin-like regions (such as C1-INH). In contrast, other glycoproteases can only target a subset of mucins. For example, TagA requires more extensive densely glycosylated regions whereas SslE/YghJ can digest major mucins such as MUC2 and MUC3, but it is inactive against mucins like CD43 and bovine submaxillary mucin (9, 23, 37, 51).

Due to low amino acid sequence conservation, it is not possible to differentiate proteases from glycoproteases solely from primary amino acid sequence. However, structural analyses revealed the presence of Ig-fold domains in all these enzymes (**Table 1**, domains in yellow). Moreover, several factors impact substrate targeting by glycoprotease, including glycan chain identity and density as well as amino acid composition (**Table 1**). Although some target motifs are known, the molecular bases for the recognition of specific target sequences remain poorly understood. Thus, even if structural analyses identify putative glycan-binding domains in a protein of interest, it is difficult to predict the specific substrates targeted by the putative glycoprotease. Together with the functional analysis, these studies define a functional class of secreted *O*-glycoproteases that mediate host-pathogen and host-commensal interactions.

## DISCUSSION

Glycan chains decorate proteins to accomplish many different functions. One important role of glycosylation is the protection of proteins against proteolytic degradation. However, there is growing evidence that bacterial pathogens and commensals have evolved specific proteases that overcome the steric impediment posed by carbohydrates, and indeed use glycans as recognition determinants to cleave glycoproteins right at the glycosylation site. In this work, we functionally characterize CpaA, a metzincin glycoprotease and T2SS secreted virulence factor of several medically relevant *Acinetobacter* strains (4, 7, 8, 12). Previous work identified blood coagulation proteins fV and fXII as targets of CpaA, suggesting a potential role in dissemination by interfering with the intrinsic coagulation pathway (4, 8, 12). Our current work expands the known targets of CpaA and indicates that CpaA is a broad-spectrum enzyme with the ability to cleave various *O*-linked human glycoproteins. Our MS analysis of proteolytic fragments resulting from glycoprotein treatment with CpaA revealed that CpaA has a consensus target sequence consisting of a Pro residue followed by a glycosylated Ser or Thr (P-S/T), which is unprecedented for bacterial glycoproteases. Unlike other secreted glycoproteases, CpaA activity is not affected by sialic acid and is not restricted to highly *O*-glycosylated proteins (mucins). Indeed, CpaA also cleaves sparsely *O*-glycosylated proteins, such as fetuin. Although broad-spectrum secreted or surface-exposed glycoproteases appear to be widespread in bacteria, their identification cannot be assigned based on sequence homology, and biochemical/structural analyses are required to designate them as glycoproteases.

CpaA is composed of four very similar Ig-like domains and a catalytic domain (13). The catalytic domain located at the C-terminus of CpaA exhibits all the canonical structural features of the metzincin superfamily. The four Ig-like domains are arranged in tandem and they resemble the INS domain of StcE, secreted by EHEC (13). These observations prompted us to further characterize CpaA activity. StcE specific motif, S/T*-X-S/T (the * denotes the glycosylation site) diverges from the P-S/T* motif recognized by CpaA (25). It is intriguing that despite recognizing different motifs, StcE and CpaA share substrates C1INH and CD55. Considering the different domain organization between both proteases, it is not surprising than both proteins are classified into different metzincin subfamilies and target different proteins.

We previously showed that the substrate-binding cleft of CpaA is formed by residues from its four Ig-like domains and its catalytic domain (13). Thus, to be recognized by CpaA, the glycosylated substrate has to expose the P-S/T* peptide bond targeted for hydrolysis. It has been proposed that the interaction of the *O*-glycans with residues in the Ig-like domains (referred as G sites by Noach *et al*) aid to position the targeted peptide-bond in the correct conformation to interact with the amino acids involved in catalysis (13, 16). Here, we show that CpaA activity is unaffected by the presence of sialic acid, indicating that the sialic acid can be accommodated inside the cleft but is not required for glycopeptide recognition. Our molecular modeling of CpaA with glycosylated substrates showed that linear glycans modified by sialic acid project this moiety away from the active site and into solvent, which supports our *in vitro* findings. Our modeling also revealed a possible interaction between the indole ring of W493 and the ring of the Pro residue in the targeted sequence. Although the digestion of glycoproteins with W493 mutants displayed substrate-specific behaviors, overall, the mutants were less active and, in some cases, exhibited a shift in glycosite preference. We propose that CpaA selectivity may, at least in part, be the result of W493 forming a potential H-pi interaction with the prolyl ring, minimizing the prolyl residue’s exposure to solvent, and/or sterically holding the substrate in the active site. Further structural and biochemical studies are required to uncover the structural features that enable CpaA to target such a remarkably high range of O-linked glycoproteins.

CpaA expression and secretion occurs across several medically relevant *Acinetobacter* strains. Deletion of CpaA resulted in attenuation of *A. nosocomialis* M2 virulence in a respiratory murine infection model, playing a role in the dissemination from the lungs to the spleen (8). Previously, the only known substrates for CpaA were fV and fXII, proteins involved in blood coagulation. By digesting these proteins, CpaA increases clotting time of human plasma (12, 15). We now showed that CpaA can indeed cleave multiple proteins *in vitro*. Among these are several proteins involved in regulation of complement activation, including CD55 and CD46. However, only CD55 was removed from the cell surfaces while CD46 remaining unaltered during the *A. nosocomialis* infection assay. We hypothesize that CD46 interacts with another protein that blocks CpaA access. Thus, not all CpaA targets identified by our *in vitro* experiments are representative bona fide targets of CpaA *in vivo.* An additional role of CD55 is to act as an anti-adhesive molecule that regulates the release of neutrophils (52). Degradation of CD55 increases the retention of the neutrophils to the apical epithelial surface with the concomitant reduction of the amount of neutrophils that cross the epithelium (48, 49). Nonetheless, CpaA expression and secretion is conserved across several medically-relevant *Acinetobacter* strains isolated from diverse anatomical sites (10). Considering the broad-spectrum activity of CpaA and the abundance of glycosylated proteins in the human host, we propose that the physiological role of CpaA likely extends beyond interfering with the coagulation pathway or complement cascade. Further work will be required to understand the full extent of host immunomodulation by CpaA.

Host mucins and *O*-glycoproteins are major components of mucus, and they are ubiquitously expressed cellular surfaces, where they act as physical barriers, receptor ligands, and mediators of intracellular signaling (53, 54). *O*-glycoproteins and mucins lack a consensus sequence for *O*-glycosylation, and their *O*-linked glycans are highly heterogeneous in their glycan composition, number of residues, and their linkages. However, aberrant mucin expression and glycosylation is also linked to various diseased states, making mucins reliable biomarkers (54). For example, the mucin MUC1 is aberrantly expressed in the majority of cancers diagnosed each year in the U.S. (53). Thus, the assessment of the mucin glycosylation status has high relevance for diagnosis of cancer and other diseases. Mucin domains are resistant to most commercially available proteases, which makes them difficult to analyze by traditional MS strategies. Bacterial glycoproteases have recently gained attention as tools for proteomic analysis of human glycoproteins (16, 22, 55). Our study shows that the broad-spectrum *O*-linked glycoprotease activity of CpaA is not affected by sialic acids. Moreover, it consistently digests any *O*-linked glycoprotein containing the P-S/T* sequence. These properties not only make CpaA a versatile enzyme modulating host-pathogen interactions, but also highlight it as a robust and attractive new component of the glycoprotemic toolbox.

## MATERIALS AND METHODS

### Strains, plasmids, and growth conditions

Bacterial strains and plasmids used in this study can be found in table S1. *E. coli* Stellar and *A. nosocomialis* M2 cells were grown in Lennox broth (LB) at 37°C. pWH1266-based plasmids were selected with tetracycline (5μg/ml).

### Generation of CpaA point mutations

The PCR primers used for site-directed mutagenesis are listed in table S1. To generate CpaAW493 variants, pWH-*cpaA-his-cpaB* was used as template. PCR was carried out using Phusion DNA Polymerase (Thermo Scientific). Site-directed mutagenesis was performed according to the method described by Fisher and Pei (56). Reactions were transformed into *E. coli* Stellar cells and transformants were selected on LB-agar supplemented with tetracycline. Mutagenesis was confirmed by DNA sequencing. The plasmids were then electroporated into electrocompetent *A. nosocomialis* M2 Δ*cpaAB::frt* strain.

### Protein purification

CpaA-his and its point mutation variants were purified from the supernatant of *A. nosocomialis* M2 (8). Briefly, *A. nosocomialis* M2 Δ*cpaAB::frt* carrying the pWH-based plasmids were grown in LB to mid log-log phase. CpaA-his tagged variants were purified by nickel affinity chromatography from cell-free filtered spent media as described before (8). Briefly, cells were pelleted at 8,000 × g for 10 minutes. Cell-free supernatants were obtained by filtration using Nalgene Rapid-Flow filter unit (pore size 0.2μm) (Thermo scientific), followed by concentration by approximately 3-fold using Amicon Ultra-15 (10,000 NMWL). 10X Binding buffer was added to the cell-free supernatant prior loaded onto a Ni-NTA agarose column (Gold Bio, St. Louis, MO) equilibrated with 1X binding buffer (50mM NaH_2_PO_4_, 300mM NaCl, 10mM imidazole, pH 8.0). The column was washed with 20 column volumes (CV) of Binding buffer and 10CV of Washing buffer (50mM NaH_2_PO_4_, 300mM NaCl, 50mM imidazole, pH 8.0). Proteins were eluted with Elution buffer (50mM NaH_2_PO_4_, 300mM NaCl, 300mM imidazole, pH 8.0). The purified proteins were concentrated, and buffer exchanged into 20mM HEPES, 150mM NaCl, 50% glycerol, pH 7.4 using Amicon ultra centrifugal filter units.

### CpaA Activity assay

Purified CpaA-his tagged variants (60μg/ml) were assayed in reaction buffer (20mM HEPES pH 7.4, 150mM NaCl, 1 mM ZnCl2) containing the substrates proteins (200μg/ml) in a final volume of 15μl. All reactions were incubated at 37°C overnight. Reactions were monitored by SDS-PAGE stained with Coomassie blue or InstantBlue. Substrate proteins used in this study CD55, CD46, TIM1, TIM4 C1-INH, and Erythropoietin (Sino Biological), Asialofetuin (Sigma), Fetuin and RNaseB (New England Biolabs), Etanercept (Enbrel), and Abatacept (Orencia).

### Immunoblotting

Bacterial whole cell and supernatant samples were prepared as before (7, 8). Briefly, cultures were grown to 0.5 OD600/ml, and 0.5 OD was pelleted by centrifugation and resuspended in 50μl of Laemmli buffer for the whole cell samples. Supernatant samples were obtained by TCA-precipitation of cell-free supernatants (7, 8). Protein samples were analyzed by SDS-PAGE and transferred to a nitrocellulose membrane and probed with either polyclonal anti-CpaA (1;1,000,(8)), and/or monoclonal anti-RNA polymerase (1:1,2000, Biolegend). Western blots were probed with IRDye-conjugated secondary antibodies and visualized with Odyssey CLx imagining system (Li-COR Biosciences, Lincoln, NE).

### Fetuin deglycosylation

Fetuin was deglycosylated under denaturing conditions using Protein Deglycosylation Mix II kit (New England Biolabs) according to the manufacturer’s protocol. Briefly, 100μg of Fetuin were incubated in Deglycosylation Mix Buffer 2 at 75°C for 10 min. After cooling down, the Protein Deglycosylation Mix II was added, with the exception of one tube to be used as positive control for CpaA’s activity. The glycosylated and deglycosylated fetuin proteins were then used as substrates to test CpaA’s activity as described above.

### Glycan Array

The glycan-array binding analysis was performed at the Consortium of Functional Glycomics (CFG). Two concentrations of purified CpaA (CFG_request_3530) and CpaA_E520A_ (CFG_request_3617) were screened with Version 5.4 of the CFG printed array consists of 585 glycans in replicates of 6, according to the standard procedure of the CFG.

### Flow cytometry

HeLa cells were maintained in DMEM with high glucose supplemented with 10% v/v heat-inactivated FBS (Gibco) at 37 °C, 5% CO2. HEK-293 cells were maintained in EMEM with 10% v/v heat-inactivated FBS (Gibco) at 37 °C, 5% CO2. Cells were seeded in 60 mm plates at a density of 1 × 106 cells per plate a day prior to each experiment. For infection assays, freshly transformed *A. nosocomialis* M2 cells with plasmids pWH-*cpaA-his-cpaB* and pMFH32 were incubated overnight in LB supplemented with tetracycline. Stationary phase cultures were normalized to OD 1 and washed three times with PBS before addition to cells at the indicated multiplicity of infection (MOI) in the respective cell media. Infected cells were incubated for 2.5 h at 37 °C, 5% CO2. Alternatively, cells were treated with 2 or 5 μg/ml of purified CpaA or CpaA_E520A_, as indicated, and incubated for 1 h at 37 °C, 5% CO2. Following infection or treatment with purified protein, eukaryotic cells were washed with PBS (twice) and harvested using 0.05% Trypsin (Corning). Cells were blocked with 2% v/v FBS in PBS and labeled with fluorophore-conjugated primary antibodies against CD55 (PE, Sino Biological), and CD46 (APC, Invitrogen). All incubations were for 45 min on ice and washed three times with PBS. Labeled cells were fixed with 4% v/v paraformaldehyde in PBS for 10 min, washed with PBS, resuspended in PBS to a cell concentration of 1×10^6^ cell/ml, and filtered prior to analysis. Flow cytometry was performed on a three-laser, 8-color FACSCantoII cytometer (BD Biosciences). Experiments were performed in technical triplicates with 10,000 events analyzed per sample. Data were analyzed using FlowJoTM 10.6 (FlowJo LLC) and individual values from three independent experiments (each experiment has at least three technical replicates) were combined and analyzed by Two-Way ANOVA using GraphPad Prism.

### Tryptic digest of gel-separated protein bands

CpaA digested proteins were separated using SDS-PAGE, fixed and visualized with Coomassie or InstantBlue, as described above. Bands of interest were excised and processed as previously described (37). Briefly gel bands were first destained in a solution of 50 mM NH_4_HCO_3_ / 50% ethanol for 20 minutes at room temperature with shaking at 750 rpm. Destained bands were dehydrated with 100% ethanol, vacuum-dried for 20 minutes and then rehydrated in 50 mM NH_4_HCO_3_ plus 10 mM DTT. Protein bands were reduced for 60 minutes at 56 °C with shaking then washed twice in 100% ethanol for 10 minutes to remove residual DTT. Reduced ethanol washed samples were sequentially alkylated with 55 mM Iodoacetamide in 50 mM NH_4_HCO_3_ in the dark for 45 minutes at room temperature. Alkylated samples were then washed with 50 mM NH_4_HCO_3_ followed by 100% ethanol twice for 5 minutes to remove residual Iodoacetamide then vacuum-dried. Alkylated samples were then rehydrated with 12 ng/μl trypsin (Promega) in 40 mM NH_4_HCO_3_ at 4 °C for 1 hour. Excess trypsin was removed, gel pieces were covered in 40 mM NH_4_HCO_3_ and incubated overnight at 37 °C. Peptides were concentrated and desalted using C18 stage tips (57, 58) before analysis by LC-MS.

### Identification CpaA digestion products and cleavage sites using reversed phase LC-MS

Purified peptides prepared were re-suspend in Buffer A* (0.1% TFA, 2% acetonitrile)and separated using a two-column chromatography set up composed of a PepMap100 C18 20 mm × 75 μm trap and a PepMap C18 500 mm × 75 μm analytical column (Thermo Fisher Scientific). Samples were concentrated onto the trap column at 5 μL/min for 5 minutes with Buffer A (0.1% formic acid, 2% DMSO) and infused into a Orbitrap Fusion™ Lumos™ Tribrid™ Mass Spectrometer (Thermo Fisher Scientific) at 300 nl/minute via the analytical column using a Dionex Ultimate 3000 UPLC (Thermo Fisher Scientific). 45 or 65 minute gradients were run for each sample altering the buffer composition from 1% buffer B (0.1% formic acid, 77.9% acetonitrile, 2% DMSO) to 28% B over 20 or 40 minutes, then from 28% B to 40% B over 5 minutes, then from 40% B to 100% B over 2 minutes, the composition was held at 100% B for 3 minutes, and then dropped to 3% B over 5 minutes and held at 3% B for another 10 minutes. For 45-minute gradients the Lumos™ Mass Spectrometer was operated in a data-dependent mode automatically switching between the acquisition of a single Orbitrap MS scan (240,000 resolution) every 3 seconds and MS2 events. For each ion selected CID (FTMS, 15K resolution, maximum fill time 100 ms, AGC 2*10^5^); HCD (FTMS, 15K resolution, maximum fill time 120 ms, normalize collision energry 35, AGC 2*10^5^); EThcD (FTMS, 15K resolution, maximum fill time 120 ms, supplementary activation 15%, AGC 2*10^5^). For 65 minute gradients the Lumos™ Mass Spectrometer was operated in a data-dependent mode automatically switching between the acquisition of a single Orbitrap MS scan (120,000 resolution) every 3 seconds and MS2 HCD scans of precursors (NCE 30%, maximal injection time of 22 ms, AGC 5*10^4^ with a resolution of 15000). HexNAc oxonium ion (204.087 m/z) product-dependent MS/MS analysis (PMID: 22701174) was used to trigger two additional scans of potential glycopeptides; a CID (ITMS, maximum fill time 100 ms, AGC 5*10^4^) scan and a EThcD (FTMS, 30K resolution, maximum fill time 200 ms, supplementary activation 15%, AGC 2*10^5^) scan.

### Mass spectrometry data analysis

The assessment of the protein coverage within CpaA digested band and the identification of Mucin glycopeptides was accomplished using MaxQuant (v1.5.3.30) (59). Searches were performed against the custom databases populated with the protein sequence of the recombinant proteins of interests with carbamidomethylation of cysteine set as a fixed modification. Searches were performed with semi-trypsin cleavage specificity allowing 2 miscleavage events. For the identification of glycopeptide multiple searches were performed on each sample allowing oxidation of methionine and a maximum of three glycan variable modifications using HexNAc (ST), HexHexNAc (ST), HexNAcHexNAc (ST) or HexNAc (ST), Hex(1)HexNAc(1)NeuAc(1) (ST), Hex(2)HexNAc(1)NeuAc(1) (ST). The precursor mass tolerance was set to 20 parts-per-million (ppm) for the first search and 10 ppm for the main search, with a maximum false discovery rate (FDR) of 1.0% set for protein and peptide identifications. The resulting protein group output was processed within the Perseus (v1.4.0.6)(59) analysis environment to remove reverse matches and common protein contaminates prior. Semi-tryptic peptide and semi-tryptic glycopeptides were manually inspected for correctness. Annotation of MSMS provided within the supplementary data was undertaken using the Interactive Peptide Spectral Annotator (59). The mass spectrometry proteomics data have been deposited to the ProteomeXchange Consortium via the PRIDE (60) partner repository with the dataset identifier PXD019941.

### Molecular modeling

All computational work was performed using the 2019 Molecular Operating Environment (MOE 2019.01) software suite available from Chemical Computing Group ULC (Montreal, Canada) (61). The X-ray crystal structures of the metzincin enzymes CpaA (6O38, (13) and serralysin (3VI1) as well as the gluzincin enzymes IMPa (5KDX, (16), ZmpB (5KDU and 5KDS, (16)), and BT4244 (5KD8, (16) were overlaid using the residues of the highly-conserved alpha helix within their active sites (Table S2).

Following a previous protocol (22), the crystal structure of CpaA was prepared by adding unresolved side chains and hydrogens as well as capping termini with acetyl or *N*Me goups. Notably, the crystallographic chaperone protein CpaB was not included during this study. This protein arranges its C-terminal tail into the CpaA catalytic site similar to zymogens of related metallopeptidases (62), and we assume that this portion of CpaB is displaced by substrate.

Peptidomimetic/peptidic ligands have been observed to bind in a similar conformation within the metzincin active site (44, 45), using the P2-P1’ ligand residues to form an antiparallel β-sheet with a β-strand of the enzyme active site. Similar to previous work (22), the crystallographic peptide ligand bound to serralysin (RPKPQQ) was used as a scaffold to construct the peptide portion (Ac-EAPSA-*N*Me) of the fetuin substrate fragment bound to the CpaA model (Table S2). Related gluzincin enzymes were found to have glycan-containing ligands near the P1’ site of their enzyme active sites (16). Thus, the crystallographic glycan bound to IMPa (Galβ1-3GalNAcα1-, 5KDX, (16) and ZmpB (Galβ1-3(Neu5Acα2-6)GalNAcα1-, 5KDU, (16)) were grafted onto the fetuin peptide fragment to generate corresponding glycoforms, and the Glycan Fragment Database (63) was used to identify a glycan (Neu5Acα2-3Galβ1-3GalNAcα1-, 2CWG, (64)) to generate the final glycoform (Table S2).

The glycopeptides were docked into the CpaA model in three steps: conformational search, virtual screen, and minimization. This process allowed for exploration of all reasonable conformations of each glycopeptide prior to induced fit docking with the CpaA model. Each glycopeptide underwent conformational search using the Amber10:EHT force field (65) to generate a corresponding library of conformers. During this step, the side chains of the peptide and the pendant groups of each sugar were allowed to freely rotate; any sialic acid moieties were also allowed to move freely. Otherwise, all other atoms were held fixed. This process generated small libraries of approximately 4,000 conformers for each glycopeptide species. The conformers of each glycopeptide library underwent virtual screening with the CpaA model, again using the Amber10:EHT force field. During this process, all atoms of the enzyme and each glycopeptide were held fixed, and the docking score for each resulting complex was calculated using the GBVI/WSA dG scoring function (66). Finally, the top ten complexes identified from the virtual screening were subsequently minimized using the Amber10:EHT force field. The glycopeptide substrate, the catalytic zinc ion, and those residues of the CpaA model having atoms within 10 Å of the substrate were allowed to move; all other residues were held fixed and solvent molecules were omitted. The best, most consistent complexes are shown.

This screening and minimization process was repeated for the Galβ1-3GalNAcα1-modified glycopeptide conformer library with the mutant CpaA models CpaAW493F, CpaAW493L, and CpaAW493A. In all cases, the dihedral angles of the peptide backbones (φ, ψ) and side chains (χ) as well as the initial glycosidic linkages (φ, ψ) of the docked substrates were measured to ensure proper geometry.

## AKNOWLEDGEMENTS

Flow cytometry experiments were performed at the Flow Cytometry & Fluorescence Activated Cell Sorting Core (Department of Pathology & Immunology, Washington University School of Medicine in St. Louis). Protein-Glycan Interaction Resource of the CFG (supporting grant R24 GM098791) and the National Center for Functional Glycomics (NCFG) at Beth Israel Deaconess Medical Center, Harvard Medical School (supporting grant P41 GM103694). We thank the Melbourne Mass Spectrometry and Proteomics Facility of The Bio21 Molecular Science and Biotechnology Institute for access to MS instrumentation. This work was supported by a National Health and Medical Research Council of Australia (NHMRC) project grant awarded to N.E.S (APP1100164) and a Canadian Institutes of Health Research project grant to ABB (PJT 159786). We thank to Dr. Paula Magnelli (New England Biolabs) for generously providing us Etanercept and Abatacept. We thank the members of the Feldman lab for critical reading of the manuscript. This work was supported by grants from the National Institute of Allergy and Infectious Diseases grant R01AI144120 awarded to M.F.F.

## AUTHOR CONTRIBUTIONS

Conceptualization: M.F.H., M.F.F. Methodology: M.F.H., N.E.S., G.DV., J.L., B.P., A.B.B., M.J.F.; Analysis: M.F.H., N.E.S., M.J.F., M.F.F.; Writing – Original Draft: M.F.H., M.F.F., Writing – Review & Editing: M.F.H., N.E.S.,G.DV., J.L. A.B.B., M.J.F., M.F.F.; Supervision: M.F.F.

## REFERENCES

1. Tacconelli, E., Carrara, E., Savoldi, A., Kattula, D., and Burkert, F. 2013. Global priority list of antibiotic-resistant bacteria to guide research, discovery, and development of new antibiotics.World Health Organization.

2. Giammanco, A, Calà C, Fasciana, T, Dowzicky, MJ. 2017. Global Assessment of the Activity of Tigecycline against Multidrug-Resistant Gram-Negative Pathogens between 2004 and 2014 as Part of the Tigecycline Evaluation and Surveillance Trial. mSphere 2.

3. Elhosseiny, NM, Elhezawy, NB, Attia, AS. 2019. Comparative proteomics analyses of Acinetobacter baumannii strains ATCC 17978 and AB5075 reveal the differential role of type II secretion system secretomes in lung colonization and ciprofloxacin resistance. Microb Pathog 128:20–27.

4. Waack, U, Johnson, TL, Chedid, K, Xi, C, Simmons, LA, Mobley, HLT, Sandkvist, M. 2017. Targeting the type II secretion system: Development, optimization, and validation of a high-throughput screen for the identification of small molecule inhibitors. Front Cell Infect Microbiol 7:380.

5. Weber, BS, Kinsella, RL, Harding, CM, Feldman, MF. 2017. The Secrets of Acinetobacter Secretion. Trends Microbiol.

6. Elhosseiny, NM, El-Tayeb, OM, Yassin, AS, Lory, S, Attia, AS. 2016. The secretome of Acinetobacter baumannii ATCC 17978 type II secretion system reveals a novel plasmid encoded phospholipase that could be implicated in lung colonization. Int J Med Microbiol 306:633–641.

7. Harding, CM, Kinsella, RL, Palmer, LD, Skaar, EP, Feldman, MF. 2016. Medically Relevant Acinetobacter Species Require a Type II Secretion System and Specific Membrane-Associated Chaperones for the Export of Multiple Substrates and Full Virulence. PLoS Pathog 12:e1005391.

8. Kinsella, RL, Lopez, J, Palmer, LD, Salinas, ND, Skaar, EP, Tolia, NH, Feldman, MF. 2017. Defining the interaction of the protease CpaA with its type II secretion chaperone CpaB and its contribution to virulence in Acinetobacter species. J Biol Chem 292:19628–19638.

9. Eijkelkamp, BA, Stroeher, UH, Hassan, KA, Paulsen, IT, Brown, MH. 2014. Comparative analysis of surface-exposed virulence factors of Acinetobacter baumannii. BMC Genomics 15:1020.

10. Wang, N, Ozer, EA, Mandel, MJ, Hauser, AR. 2014. Genome-wide identification of Acinetobacter baumannii genes necessary for persistence in the lung. MBio 5:e01163–14.

11. Johnson, TL, Waack, U, Smith, S, Mobley, H, Sandkvist, M. 2016. Acinetobacter baumannii is dependent on the type II secretion system and its substrate LipA for lipid utilization and in vivo fitness. J Bacteriol 198:711–719.

12. Tilley, D, Law, R, Warren, S, Samis, JA, Kumar, A. 2014. CpaA a novel protease from Acinetobacter baumannii clinical isolates deregulates blood coagulation. FEMS Microbiol Lett 356:53–61.

13. Urusova D V., Kinsella, RL, Salinas, ND, Haurat, MF, Feldman, MF, Tolia, NH. 2019. The structure of Acinetobacter-secreted protease CpaA complexed with its chaperone CpaB reveals a novel mode of a T2SS chaperone-substrate interaction. J Biol Chem 294:13344–13354.

14. Rosenau, F, Tommassen, J, Jaeger, KE. 2004. Lipase-specific, foldases, p. 152–161. In ChemBioChem.

15. Waack, U, Warnock, M, Yee, A, Huttinger, Z, Smith, S, Kumar, A, Deroux, A, Ginsburg, D, Mobley, HLT, Lawrence, DA, Sandkvist, M. 2018. CpaA Is a Glycan-Specific Adamalysin-like Protease Secreted by Acinetobacter baumannii That Inactivates Coagulation Factor XII. MBio 9.

16. Noach, I, Ficko-Blean, E, Pluvinage, B, Stuart, C, Jenkins, ML, Brochu, D, Buenbrazo, N, Wakarchuk, W, Burke, JE, Gilbert, M, Boraston, AB. 2017. Recognition of protein-linked glycans as a determinant of peptidase activity. Proc Natl Acad Sci U S A 114:E679–E688.

17. Yu, ACY, Worrall, LJ, Strynadka, NCJ. 2012. Structural insight into the bacterial mucinase StcE essential to adhesion and immune evasion during enterohemorrhagic E. coli infection. Structure 20:707–717.

18. Cerdà-Costa, N, Gomis-Rüth, FX. 2014. Architecture and function of metallopeptidase catalytic domains. Protein Sci. Cold Spring Harbor Laboratory Press.

19. Gomiz-Rüth, FX. 2009. Catalytic domain architecture of metzincin metalloproteases. J Biol Chem.

20. Lathem, WW, Grys, TE, Witowski, SE, Torres, AG, Kaper, JB, Tarr, PI, Welch, RA. 2002. StcE, a metalloprotease secreted by Escherichia coli O157:H7, specifically cleaves C1 esterase inhibitor. Mol Microbiol 45:277–288.

21. Nakjang, S, Ndeh, DA, Wipat, A, Bolam, DN, Hirt, RP. 2012. A novel extracellular metallopeptidase domain shared by animal Host-Associated mutualistic and pathogenic microbes. PLoS One 7:e30287.

22. Malaker, SA, Pedram, K, Ferracane, MJ, Bensing, BA, Krishnan, V, Pett, C, Yu, J, Woods, EC, Kramer, JR, Westerlind, U, Dorigo, O, Bertozzi, CR. 2019. The mucin-selective protease StcE enables molecular and functional analysis of human cancer-associated mucins. Proc Natl Acad Sci U S A 116:7278–7287.

23. Szabady, RL, Welch, RA. 2013. StcE Peptidase and the StcE-Like, Metalloendopeptidases, p. 1272–1280. In Handbook of Proteolytic Enzymes.

24. Hang, HC, Bertozzi, CR. 2005. The chemistry and biology of mucin-type O-linked glycosylation. Bioorg Med Chem 13:5021–5034.

25. Kesimer, M, Sheehan, JK. 2012. Mass spectrometric analysis of mucin core proteins. Methods Mol Biol 842:67–79.

26. Szabady, RL, Yanta, JH, Halladin, DK, Schofield, MJ, Welch, RA. 2011. TagA is a secreted protease of Vibrio cholerae that specifically cleaves mucin glycoproteins. Microbiology 157:516–525.

27. Grys, TE, Siegel, MB, Lathem, WW, Welch, RA. 2005. The StcE protease contributes to intimate adherence of enterohemorrhagic Escherichia coli O157:H7 to host cells. Infect Immun 73:1295–1303.

28. Nesta, B, Valeri, M, Spagnuolo, A, Rosini, R, Mora, M, Donato, P, Alteri, CJ, Del Vecchio, M, Buccato, S, Pezzicoli, A, Bertoldi, I, Buzzigoli, L, Tuscano, G, Falduto, M, Rippa, V, Ashhab, Y, Bensi, G, Fontana, MR, Seib, KL, Mobley, HLT, Pizza, M, Soriani, M, Serino, L. 2014. SslE Elicits Functional Antibodies That Impair In Vitro Mucinase Activity and In Vivo Colonization by Both Intestinal and Extraintestinal Escherichia coli Strains. PLoS Pathog 10:e1004124.

29. Bardoel, BW, Hartsink, D, Vughs, MM, de Haas, CJC, van Strijp, JAG, van Kessel, KPM. 2012. Identification of an immunomodulating metalloprotease of Pseudomonas aeruginosa (IMPa). Cell Microbiol 14:902–913.

30. Peppel, K, Poltorak, A, Melhado, I, Jirik, F, Beutler, B. 1993. Expression of a TNF inhibitor in transgenic mice. J Immunol 151:5699–5703.

31. Moreland, L, Bate, G, Kirkpatrick, P. 2006. Abatacept. Nat Rev Drug Discov.

32. Peppel, K, Crawford, D, Beutler, B. 1991. A tumor necrosis factor (TNF) receptor-IgG heavy chain chimeric protein as a bivalent antagonist of TNF activity. J Exp Med 174:1483–1489.

33. Grys, TE, Walters, LL, Welch, RA. 2006. Characterization of the StcE protease activity of Escherichia coli O157:H7. J Bacteriol 188:4646–4653.

34. Varki, A. 2008. Sialic acids in human health and disease. Trends Mol Med. NIH Public Access.

35. Abdullah, KM, Udoh, EA, Shewen, PE, Mellors, A. 1992. A neutral glycoprotease of Pasteurella haemolytica A1 specifically cleaves O-sialoglycoproteins. Infect Immun 60:56–62.

36. Yang, W, Ao, M, Hu, Y, Li, QK, Zhang, H. 2018. Mapping the O-glycoproteome using site-specific extraction of O-linked glycopeptides (EXoO). Mol Syst Biol 14.

37. Musumeci, MA, Hug, I, Scott, NE, Ielmini, MV, Foster, LJ, Wang, PG, Feldman, MF. 2013. In Vitro activity of Neisseria meningitidis PglL O-oligosaccharyltransferase with diverse synthetic lipid donors and a UDP-activated sugar. J Biol Chem 288:10578–10587.

38. Crooks, GE, Hon, G, Chandonia, JM, Brenner, SE. 2004. WebLogo: A sequence logo generator. Genome Res 14:1188–1190.

39. Yang, X, Tao, S, Orlando, R, Brockhausen, I, Kan, FWK. 2012. Structures and biosynthesis of the N- and O-glycans of recombinant human oviduct-specific glycoprotein expressed in human embryonic kidney cells. Carbohydr Res 358:47–55.

40. Brockhausen, I, Stanley, P. 2017. Chapter 10 O-GalNAc Glycans. Essentials Glycobiol 1:1–9.

41. Koropatkin, N, Martens, EC, Gordon, JI, Smith, TJ. 2009. Structure of a SusD homologue, BT1043, involved in mucin O-glycan utilization in a prominent human gut symbiont. Biochemistry 48:1532–1542.

42. Uhlén, M, Fagerberg, L, Hallström, BM, Lindskog, C, Oksvold, P, Mardinoglu, A, Sivertsson Å, Kampf, C, Sjöstedt, E, Asplund, A, Olsson, IM, Edlund, K, Lundberg, E, Navani, S, Szigyarto, CAK, Odeberg, J, Djureinovic, D, Takanen, JO, Hober, S, Alm, T, Edqvist, PH, Berling, H, Tegel, H, Mulder, J, Rockberg, J, Nilsson, P, Schwenk, JM, Hamsten, M, Von Feilitzen, K, Forsberg, M, Persson, L, Johansson, F, Zwahlen, M, Von Heijne, G, Nielsen, J, Pontén, F. 2015. Tissue-based map of the human proteome. Science (80-) 347:1260419–1260419.

43. Croset, A, Delafosse, L, Gaudry, JP, Arod, C, Glez, L, Losberger, C, Begue, D, Krstanovic, A, Robert, F, Vilbois, F, Chevalet, L, Antonsson, B. 2012. Differences in the glycosylation of recombinant proteins expressed in HEK and CHO cells. J Biotechnol 161:336–348.

44. Grams, F, Dive, V, Yiotakis, A, Yiallouros, I, Vassiliou, S, Zwilling, R, Bode, W, Stocker, W. 1996. Structure of astacin with a transition-state analogue inhibitor. Nat Struct Biol 3:671–675.

45. Oberholzer, AE, Bumann, M, Hege, T, Russo, S, Baumann, U. 2009. Metzincin’s canonical methionine is responsible for the structural integrity of the zinc-binding site. Biol Chem 390:875–881.

46. Coltart, DM, Royyuru, AK, Williams, LJ, Glunz, PW, Sames, D, Kuduk, SD, Schwarz, JB, Chen, XT, Danishefsky, SJ, Live, DH. 2002. Principles of mucin architecture: Structural studies on synthetic glycopeptides bearing clustered mono-, di-, tri-, and hexasaccharide glycodomains. J Am Chem Soc 124:9833–9844.

47. Tapader, R, Basu, S, Pal, A. 2019. Secreted proteases: A new insight in the pathogenesis of extraintestinal pathogenic Escherichia coli. Int J Med Microbiol. Elsevier GmbH.

48. Furniss, CRD, Low, WW, Mavridou, DAI, Dagley, LF, Webb, AI, Tate, EW, Clements, A. 2018. Plasma membrane profiling during enterohemorrhagic E. Coli infection reveals that the metalloprotease StcE cleaves CD55 from host epithelial surfaces. J Biol Chem 293:17188–17199.

49. Szabady, RL, Lokuta, MA, Walters, KB, Huttenlocher, A, Welch, RA. 2009. Modulation of neutrophil function by a secreted mucinase of Escherichia coli O157:H7. PLoS Pathog 5.

50. Hews, CL, Tran, SL, Wegmann, U, Brett, B, Walsham, ADS, Kavanaugh, D, Ward, NJ, Juge, N, Schüller, S. 2017. The StcE metalloprotease of enterohaemorrhagic Escherichia coli reduces the inner mucus layer and promotes adherence to human colonic epithelium ex vivo. Cell Microbiol 19.

51. Shon, DJ, Malaker, S, Pedram, K, Yang, E, Krishnan, V, Dorigo, O, Bertozzi, C. 2020. An Enzymatic Toolkit for Selective, Proteolysis, Detection, and Visualization of Mucin-Domain Glycoproteins.

52. Lawrence, DW, Bruyninckx, WJ, Louis, NA, Lublin, DM, Stahl, GL, Parkos, CA, Colgan, SP. 2003. Antiadhesive role of apical decay-accelerating factor (CD55) in human neutrophil transmigration across mucosal epithelia. J Exp Med 198:999–1010.

53. Beatson, R, Tajadura-Ortega, V, Achkova, D, Picco, G, Tsourouktsoglou, TD, Klausing, S, Hillier, M, Maher, J, Noll, T, Crocker, PR, Taylor-Papadimitriou, J, Burchell, JM. 2016. The mucin MUC1 modulates the tumor immunological microenvironment through engagement of the lectin Siglec-9. Nat Immunol 17:1273–1281.

54. Pinho, SS, Reis, CA. 2015. Glycosylation in cancer: Mechanisms and clinical implications. Nat Rev Cancer. Nature Publishing Group.

55. Zipfel, PF, Skerka, C. 2009. Complement regulators and inhibitory proteins. Nat Rev Immunol.

56. Fisher, CL, Pei, GK. 1997. Modification of a PCR-based site-directed mutagenesis method. Biotechniques 23:570–574.

57. Rappsilber, J, Ishihama, Y, Mann, M. 2003. Stop And Go Extraction tips for matrix-assisted laser desorption/ionization, nanoelectrospray, and LC/MS sample pretreatment in proteomics. Anal Chem 75:663–670.

58. Rappsilber, J, Mann, M, Ishihama, Y. 2007. Protocol for micro-purification, enrichment, pre-fractionation and storage of peptides for proteomics using StageTips. Nat Protoc 2:1896–1906.

59. Cox, J, Mann, M. 2008. MaxQuant enables high peptide identification rates, individualized p.p.b.-range mass accuracies and proteome-wide protein quantification. Nat Biotechnol 26:1367–1372.

60. Perez-Riverol, Y, Csordas, A, Bai, J, Bernal-Llinares, M, Hewapathirana, S, Kundu, DJ, Inuganti, A, Griss, J, Mayer, G, Eisenacher, M, Pérez, E, Uszkoreit, J, Pfeuffer, J, Sachsenberg, T, Yilmaz Ş, Tiwary, S, Cox, J, Audain, E, Walzer, M, Jarnuczak, AF, Ternent, T, Brazma, A, Vizcaíno, JA. 2019. The PRIDE database and related tools and resources in 2019: Improving support for quantification data. Nucleic Acids Res 47:D442–D450.

61. Chemical Computing Group ULC, 1010 Sherbrooke St. West, Suite #910, Montreal, QC, Canada H 2R7. 2019. Molecular Operating Environment (MOE), 2019.01.

62. López-Pelegrín, M, Cerdà-Costa, N, Martínez-Jiménez, F, Cintas-Pedrola, A, Canals, A, Peinado, JR, Marti-Renom, MA, López-Otín, C, Arolas, JL, Gomis-Rüth, FX. 2013. A novel family of soluble minimal scaffolds provides structural insight into the catalytic domains of integral membrane metallopeptidases. J Biol Chem 288:21279–21294.

63. Jo, S, Im, W. 2013. Glycan fragment database: A database of PDB-based glycan 3D structures. Nucleic Acids Res 41.

64. Wright, CS, Jaeger, J. 1993. Crystallographic refinement and structure analysis of the complex of wheat germ agglutinin with a bivalent sialoglycopeptide from glycophorin A. J Mol Biol 232:620–638.

65. Weiner, SJ, Kollman, PA, Singh, UC, Case, DA, Ghio, C, Alagona, G, Profeta, S, Weiner, P. 1984. A New Force Field for Molecular Mechanical Simulation of Nucleic Acids and Proteins. J Am Chem Soc 106:765–784.

66. Corbeil, CR, Williams, CI, Labute, P. 2012. Variability in docking success rates due to dataset preparation. J Comput Aided Mol Des. J Comput Aided Mol Des.

67. Bardoel, BW, Hartsink, D, Vughs, MM, de Haas, CJC, van Strijp, JAG, van Kessel, KPM. 2012. Identification of an immunomodulating metalloprotease of Pseudomonas aeruginosa (IMPa). Cell Microbiol 14:902–913.

68. Seo, J, Brencic, A, Darwin, AJ. 2009. Analysis of secretin-induced stress in Pseudomonas aeruginosa suggests prevention rather than response and identifies a novel protein involved in secretin function. J Bacteriol 191:898–908.

69. Gutiérrez-Jiménez, J, Arciniega, I, Navarro-García, F. 2008. The serine protease motif of Pic mediates a dose-dependent mucolytic activity after binding to sugar constituents of the mucin substrate. Microb Pathog 45:115–123.

70. Jiang, P, Mellors, A. 2013. O-Sialoglycoprotein, Endopeptidase, p. 1664–1666. In Handbook of Proteolytic Enzymes. Elsevier Ltd.

71. Tapader, R, Bose, D, Basu, P, Mondal, M, Mondal, A, Chatterjee, NS, Dutta, P, Basu, S, Bhadra, RK, Pal, A. 2016. Role in proinflammatory response of YghJ, a secreted metalloprotease from neonatal septicemic Escherichia coli. Int J Med Microbiol 306:554–565.

72. Cladman, WM, Watt MA V., Dini, JP, Mellors, A. 1996. The pasteurella haemolytica O-sialoglycoprotein endopeptidase is inhibited by zinc ions and does not cleave fetuin. Biochem Biophys Res Commun 220:141–146.

